# High-Throughput Screening Identifies PLK1 Inhibition as a Strategy to Potentiate BTK Blockade in Marginal Zone Lymphoma

**DOI:** 10.1101/2025.10.25.684506

**Authors:** Alex Zadro, Alberto Arribas, Michela Robusto, Stefania Vultaggio, Adrian Andronache, Giovanni Faga, Kristyna Kupcova, Eleonora Cannas, Filippo Spriano, Giulio Sartori, Luciano Cascione, Chiara Tarantelli, Sara Napoli, Andrea Rinaldi, Davide Rossi, Emanuele Zucca, Matteo Moretti, Ondrej Havranek, Anastasios Stathis, Mario Varasi, Ciro Mercurio, Francesco Bertoni

## Abstract

B-cell receptor (BCR) signaling is a key therapeutic target in B-cell lymphomas, and Bruton tyrosine kinase inhibitors (BTKi) have demonstrated clinical efficacy in marginal zone lymphoma (MZL). However, the rate of complete remissions is relatively low, and resistance remains a significant clinical challenge, underscoring the need for novel combination strategies. To identify compounds that enhance the activity of BTKi and overcome resistance, we conducted a high-throughput screen of 1,695 compounds using a previously developed MZL model of acquired resistance to BTK and PI3K inhibitors derived from the Karpas1718 cell line. Thirty-three compounds showed single-agent anti-proliferative activity in the nanomolar range, both in the Karpas1718-resistant and parental cells. Based on their clinical potential in combination with BTKi, seven compounds were selected for further validation. The polo-like kinase 1 (PLK1) inhibitor rigosertib emerged as a top candidate, showing strong activity in both parental and BTKi-resistant cell lines. Combination treatment of rigosertib and BTKi zanubrutinib resulted in broad transcriptomic changes, characterized by the downregulation of pathways involved in B-cell activation, proliferation, and BCR signaling. Mechanistically, the combination reduced phosphorylation of key BCR pathway components and inhibited NF-κB nuclear translocation. These findings suggest that PLK1 inhibition can overcome BTKi resistance by further inhibiting BCR signaling through the canonical NF-κB pathway. Therefore, the dual pharmacological inhibition of BTK and PLK1 is a promising therapeutic approach for patients with MZL and warrants further preclinical and clinical investigations.

## Introduction

Marginal zone lymphoma (MZL) is an indolent malignancy of mature B cells, frequently triggered by chronic stimulation of the B cell receptor (BCR) signaling by persistent infections or autoantigens ^1-4^. Therefore, MZL largely relies on the BCR signaling pathway to survive and proliferate, and the constitutively activated NF-κB signaling is a common feature ^1,3-5^. After antigen exposure, physiological BCR signaling begins with the clustering of different BCR molecules and the consequent phosphorylation of the immunoreceptor tyrosine-based activation motif (ITAM) domains by SRC-family kinases ^6^. This process enables the recruitment of SYK, which forms a complex with CIN85 and BLK and is responsible for BTK phosphorylation ^7^. BTK-dependent phosphorylation of PLCγ2 leads to calcium mobilization by hydrolysis of the phosphatidylinositol-4,5-bisphosphate (PI(_4,5_)P_2_) into diacylglycerol (DAG) and inositol trisphosphate (IP_3_) ^5,8^. The final steps of the physiological BCR signaling consist of the phosphorylation, ubiquitination, and degradation of the IKK complex and the consequent translocation of the NF-κB transcription factors into the nucleus ^9^. Particularly, the translocation of p50/p65 heterodimer into the nucleus activates the NF-κB transcriptional program ^5,10^. On the other hand, tonic BCR signaling results in PI3K/AKT/mTOR activation, which is crucial for sustaining lymphoma survival and proliferation ^11^.

Due to its key role in B-cell lymphomagenesis, BCR signaling is an attractive therapeutic target for treating B-cell lymphomas, and several compounds targeting key proteins in its pathway have been developed ^1,2,5,11-13^. In particular, BTK and PI3K targeting have proven to be effective therapeutic strategies for treating B-cell lymphomas by inhibiting BCR signaling ^1,2,5,11-13^. As a result, numerous BTK and PI3K inhibitors (BTKi and PI3Ki) have been developed, and some have been successfully introduced into clinical settings, including for MZL patients ^2,11-14^. However, resistance to BCR-targeted agents often develops in patients ^15,16^. Combination therapy could help overcome resistance to treatment; therefore, identifying new therapeutic targets and effective drug combinations is essential.

We screened 1,695 compounds in a bona fide MZL model ^17^ and its derivative with secondary resistance to PI3Ki and BTKi ^18^, identifying various compounds with strong activity in both cellular models. Among these, the Polo-like Kinase-1 (PLK1) inhibitors were potent as single agents and increased the anti-tumor activity of BTK and PI3K inhibitors in MZL models, as well as in their resistant derivatives. Mechanistically, the dual pharmacological inhibition of PLK1 and BTK decreased BCR signaling and reduced p65 nuclear translocation.

## Methods

### Cell lines

Established human bona fide MZL cell lines (Karpas1718 and VL51 ^17,19^) and their derivatives with acquired resistance to PI3K/BTK inhibitors ^18,20^ or to PI3K inhibitors ^21^ were cultured in RPMI 1640 supplemented with 15% FBS, 2mM L-Glutamine, 1% Pen/Strep, 25mM Hepes, and were maintained at 37°C with 5% CO2. Cell line identity was authenticated by using short tandem repeat (STR) DNA fingerprinting with the Promega GenePrint 10 System kit (B9510). The cell lines were stored at −150°C, and all experiments were conducted within one to two months after thawing. Routine Mycoplasma testing was performed using the MycoStrip assay (rep-mysnc-100, InvivoGen) to confirm negativity.

### High-throughput compound screening

The library used for the high-throughput screening (HTS) contained 1695 small molecules, selected from a collection of FDA-approved drugs, agents approved by other regulatory bodies, natural products, and targeted molecules at various stages of clinical and preclinical development.

The HTS was performed on parental Karpas1718 and its derivative model with resistance to BTK and PI3K inhibitors, obtained with prolonged exposure to idelalisib, using the Microlab STAR Workstation (ML-STAR, Hamilton). In detail, 1750 cells were seeded into each well of a 384-well white flat-bottom assay plate (Greiner Bio-One, #781080), in 35 μL of complete medium. Plates were then automatically transferred into the robot-integrated incubator (Cytomat 2 C-Lin, Thermo Scientific) and stored for 24 hours. For drug treatment, molecules were initially arrayed at 10□mM in DMSO, according to the final layout, in 384-well PCR barcoded plates (Thermo Scientific, #BC-1384) and subsequently subjected to a 1:10 serial dilution of 5 points. A total of 41 molecules were disposed into a single compound plate, which also accommodates one negative (0.5% DMSO) and two internal controls at five doses (50-0.005μM idelalisib and 1-0.0001μM panobinostat-Selleck Chemicals #S1030). Some wells inside the assay plates were dedicated to panobinostat at 1 μM, used as a positive control for 100% cell death. On the day of the treatment, 3 μl of the drugs were further diluted in 72 μl of complete medium in temporary plates and then dispensed (5 μl) onto cells at final concentrations of 50, 5, 0.5, 0.05, and 0.005 μM.

Cell viability was assessed 72 hours post-treatment using CellTiter-Glo 2.0 (CTG; Promega, #G9243) by adding 40 μL of reagent to each well of the assay plates.

### Screening Data Analysis

The CTG/viability data read by the ENVISION multiplate reader went through a custom data management system that fully integrates compound library management, sample tracking, and data annotation and registration. As such, viability data for every well of all assay plates were associated with the corresponding experimental treatments and conditions on the different cell lines. Then, the annotated data underwent a standard processing and analysis pipeline for HTSATLAB, implemented in Matlab (MathWorks). The intra- and inter-plate raw data variability was removed using the Viability Percent data normalization (fixing 100% for Negative Controls and 0% for Positive Controls on each plate). The normalized data from all assay plates that passed Quality Control (i.e., robust Z-factor > 0.5) were further used to estimate the dose-response curves for each treatment on each cell line using the four-parameter logistic model. Promising compounds were selected as “highly active” (IC50<150nM in Karpas1718 parental and resistant cells), of “active non-anticancer” (150nM<IC50<5000nM), and validated in a further screen performed in triplicate.

### Dose-Response

Highly active compounds in parental and resistant cells were selected for further validation as a single agent or in combination with ibrutinib or idelalisib after 72 hours of exposure to increasing doses of molecules, followed by MTT assay. Sensitivity to single-drug treatments was evaluated using IC50 (4-parameter calculation based on log-scaled doses, PharmacoGX R package ^22^) calculations. The beneficial effect of the combinations versus the single agents was considered both as synergism according to the Chou-Talalay combination index ^23^, to the highest single agent (HSA) ^24^, to zero interaction potency (ZIP), synergy models included in the SynergyFinder R package ^25^; as well as potency and efficacy according to the MuSyC algorithm ^26^.

### Genomics

Total RNA was extracted using the All Prep DNA/RNA Mini Kit (Qiagen, Hilden, Germany) according to the manufacturer’s instructions. RNA concentration and integrity were measured with a NanoDrop ND-1000 spectrophotometer and assessed through the Bioanalyzer 2100 instrument (Agilent Technologies, Santa Clara, CA, USA). The NEBNext Ultra Directional RNA Library Prep Kit for Illumina (New England BioLabs Inc.) was used in conjunction with the NEBNext Multiplex Oligos for Illumina (New England BioLabs Inc.) and the NEBNext Poly(A) mRNA Magnetic Isolation Module for cDNA synthesis, incorporating barcode sequences. The pre-pool sequencing was performed using the NextSeq 2000 (Illumina, San Diego, CA, USA) with the P2 reagent kit V3 (100 cycles; Illumina). Samples were processed starting from stranded, single-ended 120bp-long sequencing reads.

We evaluated the quality of RNA-seq reads using FastQC (v0.11.5) ^27^ and removed low-quality reads/bases and adaptor sequences using Trimmomatic (v0.35) ^28^. The resulting high-quality trimmed reads were aligned to the human reference genome (GRCh38, version 113) using STAR ^29^, a spliced-read aligner that supports the alignment of reads spanning multiple exons. Samples were considered of good quality if more than 85% of sequencing reads aligned to the reference genome. Gene expression quantification was performed using HTSeq-count ^30^ with Ensembl gene annotation for GRCh38.113 as reference. Expression values were provided in tab-delimited format. We filtered the dataset to retain genes with counts-per-million (CPM) values greater than 5 in at least half of the samples. The data were normalized using the TMM (trimmed mean of M-values) method from the edgeR package ^31,32^ and transformed to log2 CPM using the cpm() function. Differential gene expression for each comparison of interest was computed using limma, following TMM normalization and voom transformation ^33^. Functional analysis was performed on the collapsed gene symbol list using Gene Set Enrichment Analysis (GSEA) on differentially expressed genes ranked by t-statistics, with the MSigDB (Molecular Signatures Database) C2-C7 gene sets ^34,35^, Hallmark, Gene Ontology Biological Process (GOBP), and Gene Ontology Cellular Components (GOCC) as reference datasets. The top 30 by |NES| significantly (adj. p value < 0.05) differentially regulated GOCC pathways in Karpas1718 were used to perform ridge plot analysis.

The top 30 significantly (FDR < 0.05) downregulated GOBP pathways in Karpas1718, upon combination treatment with the lowest NES, were plotted using the enrichMap function to perform pathway network analysis.

Expression values will be available at the National Center for Biotechnology Information (NCBI) Gene Expression Omnibus (GEO;http://www.ncbi.nlm.nih.gov/geo) database.

### Immunoblotting

Cells were seeded in T25 flasks at a density of 1 × 10^^6^ cells/mL in a final RPMI volume of 10 mL and were treated as indicated. At the end of treatment, the collected cells were centrifuged (1200 rpm, 5min), and the pellet was washed once with 1X PBS. We performed protein extraction using the M-PER buffer (Thermo Scientific, 78501), which was enriched with a 100X Halt Protease and Phosphatase Inhibitor Cocktail (Thermo Scientific, 78440). Protein samples were then quantified using the Pierce BCA Protein Assay Kit (Thermo Scientific, 23225) according to the manufacturer’s instructions and read on a Cytation 3 instrument (BioTek). To analyze proteins via immunoblotting, equal amounts of protein (30 µg per sample) were mixed with NuPAGE LDS Sample Buffer (Invitrogen, NP0008) and heated to 95 °C for 10 min. Proteins were separated on 4–20% Mini-PROTEAN TGX Precast Protein Gels (Biorad, 4561096) and transferred to nitrocellulose membranes using the Trans-Blot Turbo Mini 0.2 µm Nitrocellulose Transfer Packs (Biorad, 1704158) and the Trans-Blot Turbo Transfer System (Biorad). After blocking in 5% powdered milk (Roth, T145.2) in 1X TBST buffer, membranes were incubated overnight with 1:1000 in 5% powdered milk diluted primary antibodies (Supplementary Table 1), followed by anti-rabbit (Cytiva Europe, NA934-1ML) or anti-mouse (Cytiva Europe, NA931) HRP-linked secondary antibodies 1:2000 in 5% powdered milk. Membranes were developed with Westar Sun (Cyagen, XLS063.0250), Antares (Cyagen, XLS142.0250), or Ultra (Cyagen, XLS075.0100) substrates and visualized with the Fusion Solo S imaging system (Vilber). Vinculin or GAPDH were used as a loading controls to normalize the protein signals obtained from band densitometry analysis in Fiji ImageJ.

### Immunofluorescence

The IbiTreat µ-Slide 18 Well (Ibidi, 81816) was pre-coated by applying poly-L-ornithine solution (Sigma, A-004-C) to each well and incubating at room temperature (RT) for 1 hour. After coating, the solution was removed, and the plates were allowed to air dry. Cells were then seeded at 1 × 10^^5^ cells per well (100 µL of a 1 × 10^^6^ cells/mL suspension) under the specified treatment conditions. After 4, 6, or 8 hours, cells were fixed with 4% paraformaldehyde (Science Services, E15710) for 15 minutes at RT, followed by three washes with DPBS with calcium and magnesium (Gibco, 14040117), each lasting 5 minutes. For permeabilization, cells were incubated in 0.3% Triton-X (Sigma, T8787) in DPBS for 10 min at RT. To block non-specific binding, 4% bovine serum albumin (BSA) + 0.1% Triton-X in DPBS was added and incubated for 1 hour at RT while shaking. After blocking, the BSA solution was removed, and primary antibodies (*Supplementary Table 1*), diluted in 4% BSA and 0.1% Triton X-100 in DPBS, were added for overnight incubation at 4°C. The next day, wells were washed three times with DPBS for 5 minutes. Secondary antibodies (*Supplementary Table 1*), diluted in 4% BSA + 0.1% Triton-X with 1:1000 DAPI (Sigma, D9542), were added and incubated for 1 hour at RT. Following secondary antibody incubation, the wells were washed three times with DPBS for 5 minutes each. Samples were then stored in Vectashield mounting medium diluted (1:4 dilution) in 80% glycerol and kept at 4°C for long-term preservation. Images were acquired using the Leica TCS SP5 confocal□microscope with an HCX PL APO lambda blue 63.0 × 1.40 oil objective. To quantify the total and nuclear p65 intensity signals, a pixel classifier was trained using an artificial neural network model at full resolution (0.36 μm/px) in QuPath software ^36^. This trained classifier has been unbiasedly used to classify and quantify all the images.

### AKT activity by Förster resonance energy transfer (FRET)

Karpas1718 parental cells were transfected with the Förster resonance energy transfer (FRET)-based AKT activity reporter Lyn-AktAR2-EV (Addgene plasmid #125199) as well as with a control “dead” unresponsive version of the reporter Lyn-AktAR2-EV-D (Addgene plasmid #125200) ^37^. Levels of FRET/AKT activity were measured using flow cytometry, and data processed using the R package fRet as previously described ^37-40^.

### Graphical representation and statistical analysis

Graphical representation and statistical analyses were performed using GraphPad Prism version 8.01 (https://www.graphpad.com), unless stated otherwise. The specific statistical tests used are detailed in the respective figure legends.

## Results

### Screening of 1,695 compounds identified clinical-grade compounds that were highly active in both parental and resistant Karpas1718 cells

To identify agents with potential to overcome resistance to BTK and PI3Kδ inhibitors in MZL, we screened a custom-assembled small-molecule library comprising 1,695 compounds. This library included FDA-approved drugs, agents approved by other regulatory bodies, natural products, and targeted molecules at various stages of clinical and preclinical development (Figure 1A, Supplementary Table 2). The compounds spanned a range of mechanistic classes, including inhibitors of DNA damage and repair, cell cycle regulators (cyclin-dependent kinases, aurora and polo kinases), epigenetic and proteasome modulators, and others.

**Figure 1.**
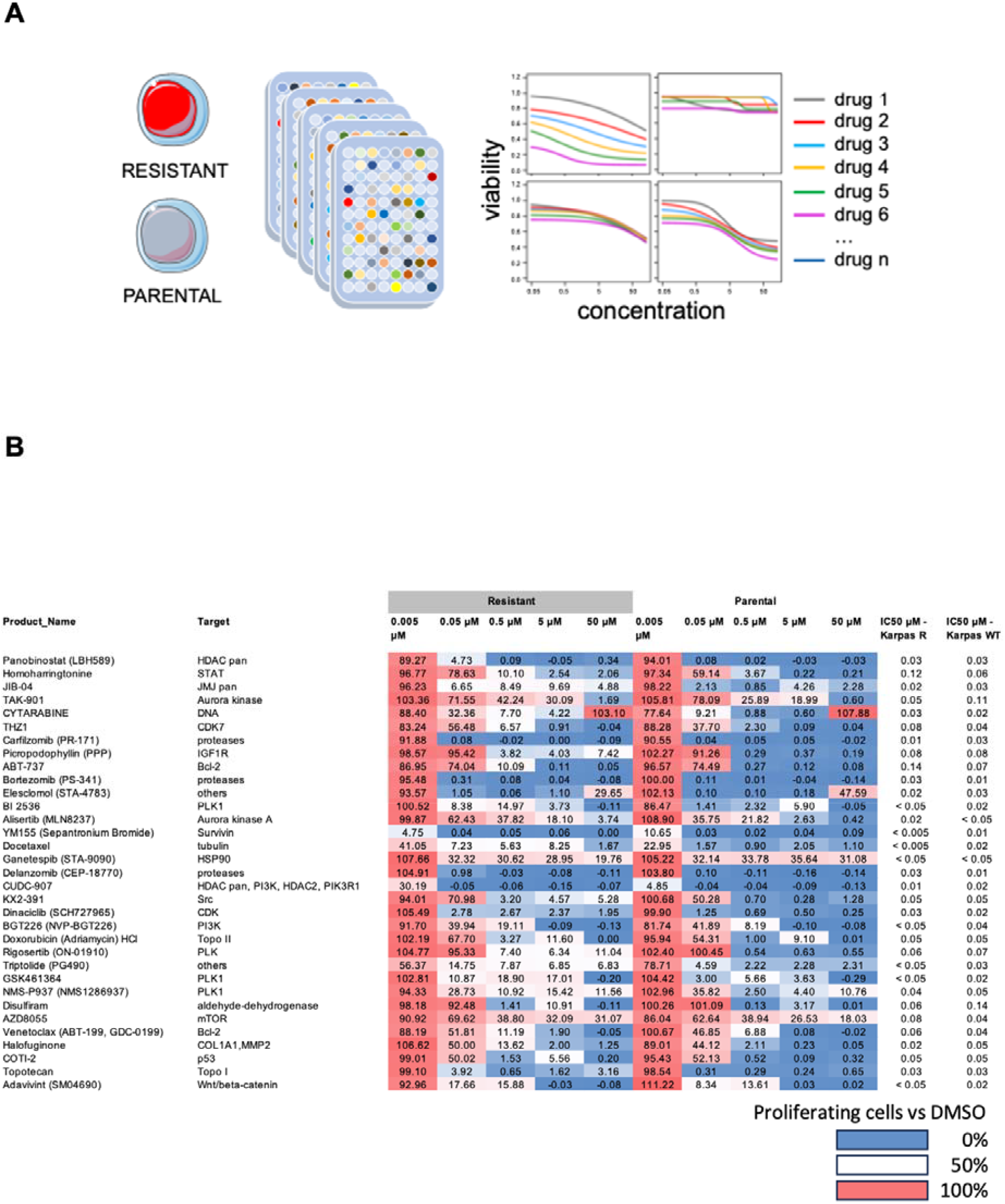
Screening of 1,695 compounds identified clinical-grade compounds that were highly active in both parental and resistant Karpas1718 cells. A) Schematic drug screening experimental design representation in parental and resistant Karpas1718 cells. B) Heatmap representing parental and resistant Karpas1718 cells proliferation upon treatment with increasing doses of the indicated compounds.

The screening was conducted in a high-throughput format using the bona fide MZL model Karpas1718 ^17^, and its derivative, which exhibits ERBB4-driven resistance to several BTK and PI3K inhibitors, obtained by prolonged exposure to the PI3Kδ inhibitor idelalisib ^18^. Each compound was tested in a 72-hour dose-response assay across a five-concentration log-range (0.005-50 µM). DMSO-treated cells served as the 100% viability control, while cells treated with the pan histone deacetylase (HDAC) inhibitor panobinostat at an established lethal dose represented the 100% cell death.

Cell viability was assessed using the CellTiter-Glo assay, which measures intracellular ATP levels as a surrogate for cell proliferation. Data were analyzed using a four-parameter logistic dose-response model.

The screen was conducted in six rounds using 384-well plates, with each plate containing internal positive (panobinostat-treated cells) and negative (DMSO-treated cells) controls. In addition, in some wells, cells were treated with idelalisib and panobinostat across a five-concentration log-range, spanning, respectively, from 0.005 to 50 µM for the former and from 0.0001 to 1 µM for the HDAC inhibitor.

Following raw data acquisition, rigorous quality control (QC) procedures were applied. Inter- and intra-plate variability was minimized through data normalization. An average Z-factor higher than 0.7 across all 384 well plates of the screening highlighted the robustness and the quality of the obtained data.

Through data analysis and visual inspection of the dose response curves, we identified 58 small molecules with an IC50 < 150 nM in both wild-type and resistant Karpas1718 cells (Supplementary Table 2). Among them, 33 small molecules (Supplementary Table 2) were selected for further investigation. These compounds target various biological processes, including apoptosis, cell cycle regulation, DNA damage response, epigenetics, cancer stem cell maintenance, and metabolic pathways (Figure 1B).

Seven clinical-grade compounds were selected for further validations based on their potential for clinical application in combination with PI3K or BTK inhibitors: rigosertib/ON-01910 (Polo-like Kinase-1 (PLK1) inhibitor), elesclomol (mitochondrial electron transport targeting agent), alisertib (Aurora Kinase A inhibitor), ganetespib (HSP90 inhibitor), fimepinostat/CUDC-907 (pan-HDAC-PI3K inhibitor), disulfiram (ALDH1 inhibitor), and panobinostat (HDAC inhibitor). The selected compounds were tested in combination with idelalisib in Karpas1718 models and in three other MZL models, the parental VL51 cells and their idelalisib and ibrutinib-resistant derivatives.

Most of the tested compounds were beneficial when combined with idelalisib, increasing the potency rather than the efficacy of the PI3K inhibitor. According to the Chou-Talalay combination index, most of the compounds were synergistic to idelalisib, including elesclomol, rigosertib, alisertib, fimepinostat, and panobinostat (Figure 2A). Disulfiram and rigosertib were selected for further experiments.

**Figure 2.**
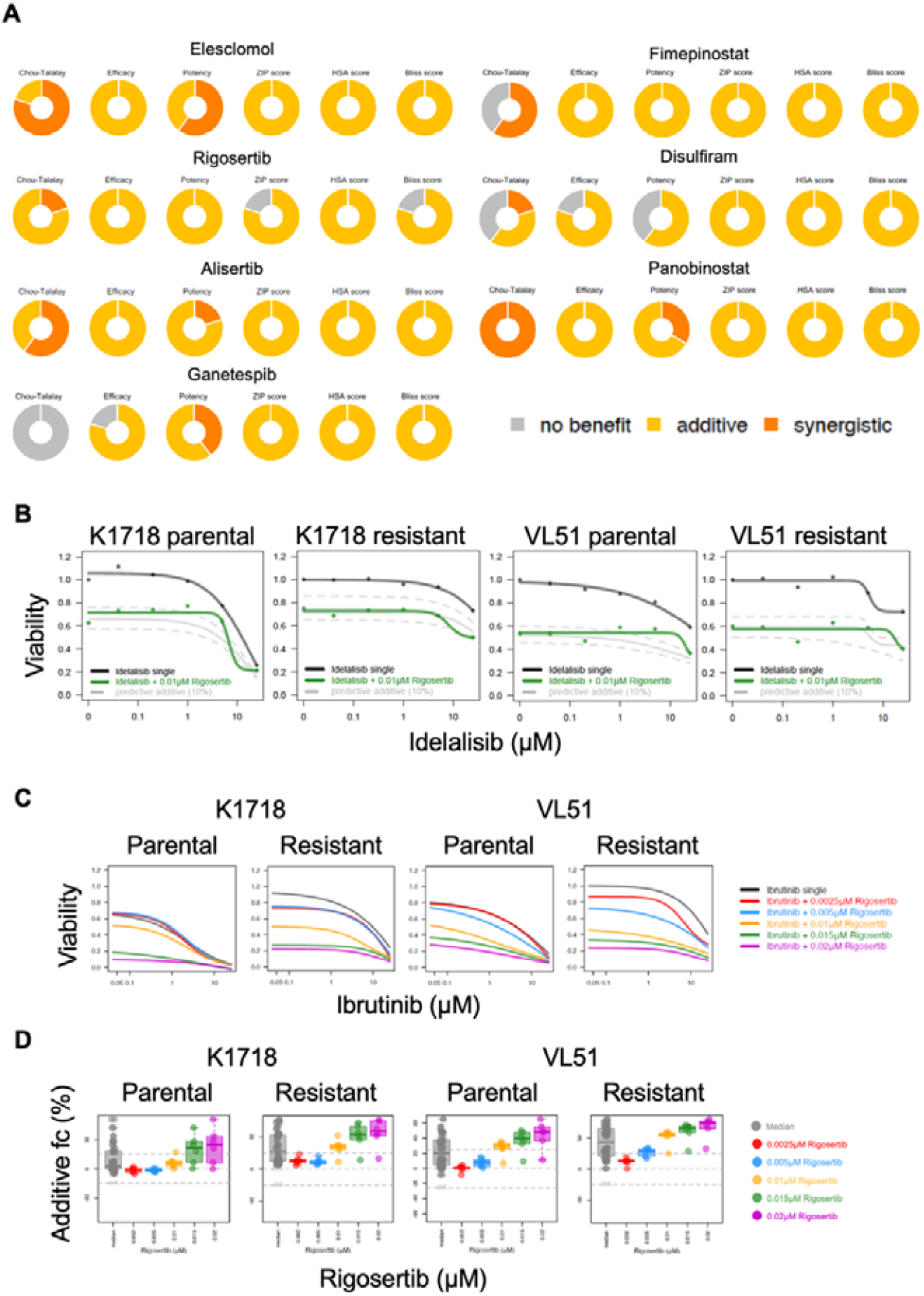
Rigosertib enhances idelalisib and ibrutinib efficacy and exhibits potent single-agent activity in parental and resistant Karpas1718 and VL51 cells. A) Synergy scores of the indicated compounds in combination with idelalisib in resistant Karpas1718 cells. Graphs generated with R software. B) Dose-response curves in parental and resistant Karpas1718 and VL51 cells treated with increasing idelalisib concentration and fixed rigosertib concentration (0.015μM). Graphs generated with R software. C) Dose-response curves in parental and resistant Karpas1718 and VL51 cells treated with increasing ibrutinib concentration and increasing rigosertib concentrations. Graphs generated with R software. D) Additive effects of increasing rigosertib concentrations to the ibrutinib combination in parental and resistant Karpas1718 and VL51 cells. Graphs generated with R software.

Disulfiram was able to overcome idelalisib resistance in Karpas1718 and VL51 PI3Ki-resistant models (Supplementary Figure 1A) and ibrutinib resistance in Karpas1718 and VL51 BTKi-resistant models (Supplementary Figure 1B).

Notably, rigosertib not only overcame resistance to idelalisib (Figure 2B) and ibrutinib (Figure 2C) in the Karpas1718 and VL51 models but also showed extensive anti-proliferative activity at increasing concentrations when combined with ibrutinib in parental cells (Figure 2C). Moreover, rigosertib demonstrated additive activity when combined with ibrutinib in all the models used (Figure 2D). Notably, three other PLK1 inhibitors (BI-2536, GSK461364, and NMS-P937) and two Aurora Kinase (AURK) inhibitors (TAK-901 and Alisertib) were among the 33 selected hits (Figure 1B), further highlighting the AURKA/PLK1 as an interesting biological axis for overcoming resistance to PI3K and BTK inhibitors.

Therefore, because of its potency as a single agent and its ability to enhance the efficacy of idelalisib and ibrutinib in both Karpas1718 and VL51 parental and resistant models, the PLK1 inhibitor rigosertib was selected for further studies.

### PLK1 inhibition downregulates pathways related to B cell proliferation and activation through reduced BCR signaling

To investigate the molecular mechanisms by which rigosertib enhances the efficacy of BTK inhibition, we used the second-generation BTK inhibitor zanubrutinib, which is an approved therapeutic option for refractory and relapsed MZL in various countries ^41^. Bulk RNA sequencing was performed on the Karpas1718 parental cell line treated with DMSO, zanubrutinib (5 nM), rigosertib (10 nM), or the zanubrutinib/rigosertib combination for 4, 8, and 12 hours.

The single treatment with zanubrutinib resulted in a robust transcriptome response, with 172 and 437 genes upregulated and downregulated, respectively. Rigosertib alone, instead, had a minor effect on the transcriptome of the cell line, with only 92 and 50 upregulated and downregulated genes, respectively. However, the combination of rigosertib and zanubrutinib resulted in a significant increase in transcriptome changes, with 889 and 906 upregulated and downregulated genes, respectively (Figure 3A). Moreover, a functional analysis of the modulated transcripts revealed that the P53 pathway was upregulated in response to zanubrutinib and rigosertib as single treatments, and this effect was even stronger when the two drugs were combined, supporting their additive effect in inducing cell apoptosis (Supplementary Figure 2A).

**Figure 3.**
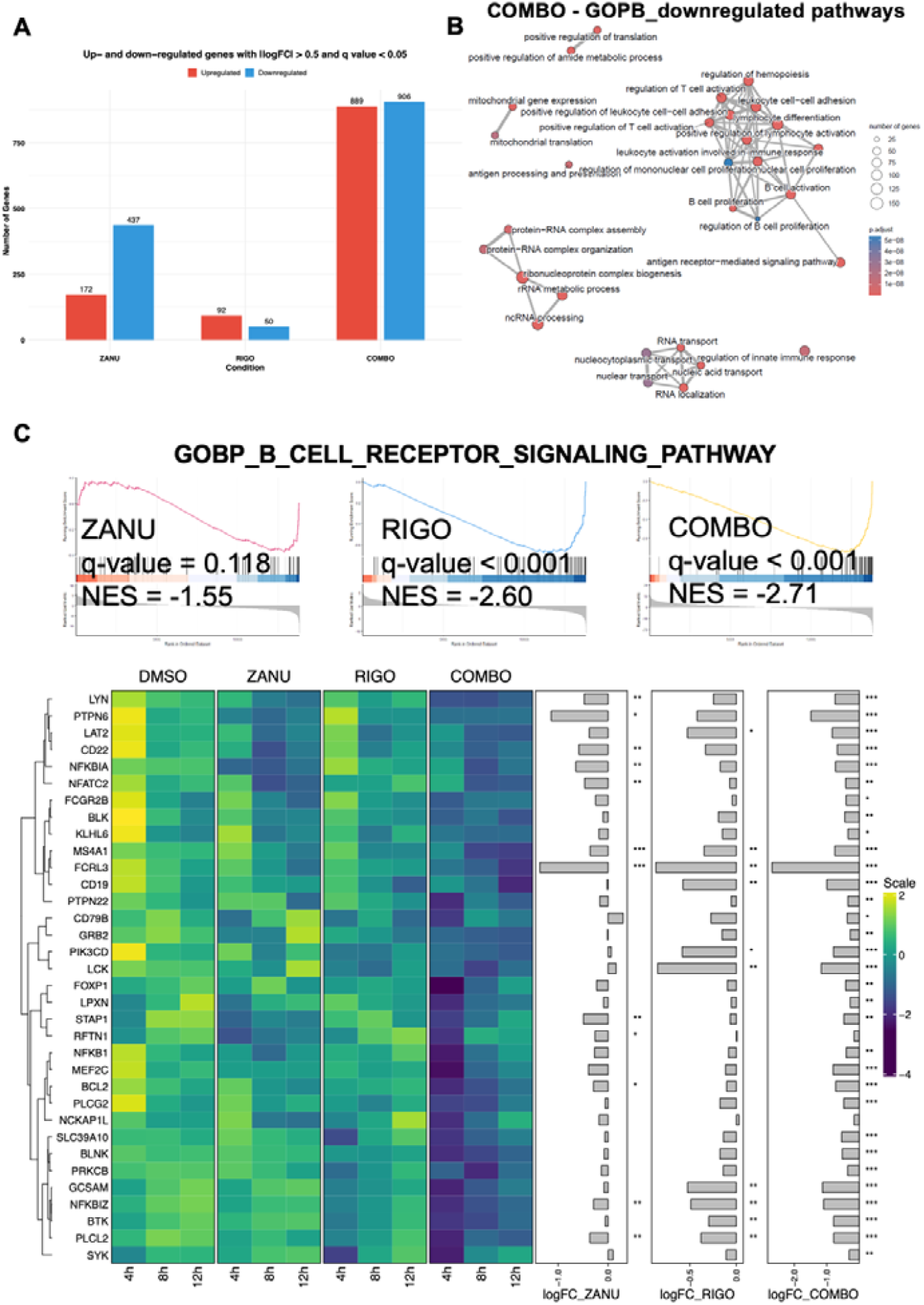
Rigosertib enhances the inhibition of the BCR signaling pathway driven by zanubrutinib. A) Number of significant (q value < 0.05) differentially expressed (|logFC| > 0.5) genes upon zanubrutinib (5nM), rigosertib (10nM) or zanubrutinib/rigosertib combination in Karpas1718 parental cells. Graph generated with R software. B) Pathway network representation of the significant GOBP pathways top-downregulated upon zanubrutinib/rigosertib combination in Karpas1718 parental cells. Graph generated with R software. C) Enrichment plots of the indicated pathway upon zanubrutinib (5nM), rigosertib (10nM) or zanubrutinib/rigosertib combination in Karpas1718 parental cells. Heatmap showing the regulation of the core enrichment genes upon the indicated treatment conditions. Graph generated with R software. Statistical significance tested with moderated t-test (*=p<0.05; **=p<0.01; ***=p<0.01). (ZANU=zanubrutinb; RIGO=rigosertib; COMBO=combination).

PLK1 is a key kinase involved in mitosis, particularly in the correct formation of the mitotic spindle by coordinating microtubule dynamics and kinetochore function ^42^. As expected, by performing ridge plot analysis on the top 30 GOCCs significantly (adj. p-value < 0.05) regulated pathways based on absolute NES, we identified several gene sets related to microtubules and the mitotic spindle that were upregulated upon rigosertib single treatment and, particularly, when rigosertib was combined with zanubrutinib (Supplementary Figure 2B). The GOCC mitotic spindle gene set was significantly upregulated by the combination (Supplementary Figure 2C). Moreover, essential genes for mitotic spindle formation, such as PLK1, AURKA, and CDK1, were upregulated at the transcript level upon the combination (Supplementary Figure 2C). This is in line with previous findings that showed upregulated PLK1 protein levels upon its pharmacological inhibition, probably related to feedback loop compensatory effects and to an accumulation of cells arrested in G2/M phase ^43^, confirming the inhibitory activity of rigosertib on PLK1 in our model.

The pathway network analysis on the most downregulated genes upon zanubrutinib and rigosertib combination identified three major clusters belonging to nuclear transport, RNA processing, and immune-related pathways of the GOBP gene sets. The latter included pro-survival signatures in B-cells, such as B-cell activation and proliferation, and BCR signaling, which were downregulated by the combination (Figure 3B). In detail, the addition of rigosertib further downregulated B-cell activation and proliferation signatures, which were already repressed by zanubrutinib alone (Supplementary Figure 3A). At the transcript level, BCL2 was significantly downregulated upon combination treatment in Karpas1718 (Supplementary Figure 3B). Additionally, we observed a trend towards a reduction in BCL2 protein levels upon treatment with zanubrutinib, rigosertib, and the combination at 48 hours (Supplementary Figure 3C-D).

As expected, zanubrutinib reduced BCR signaling pathway activity, but surprisingly, rigosertib exposure also resulted in the downregulation of this pathway. Notably, combining rigosertib with zanubrutinib further suppressed BCR signaling genes (Figure 3C). These findings suggest that the mechanism of action of the zanubrutinib and rigosertib combination might derive from a concomitant inhibition of the BCR signaling pathway.

### PLK1 inhibition enhances BTKi-driven BCR signaling inhibition through the canonical NF-κB pathway

To validate the effects of rigosertib on the BCR signaling pathway, we treated Karpas1718 parental cells with zanubrutinib (5 nM), rigosertib (10 nM), or their combination for 4, 24, and 48 hours. We analyzed the expression of three key BCR-NF-κB signaling proteins, BTK, PLCγ2, and p65, in different treatment conditions, either in total or active phosphorylated form. Total BTK protein levels remained unchanged across all treatment conditions and time points. At the same time, BTK-Y223 phosphorylation was significantly inhibited by zanubrutinib at 24 and 48 hours, an effect that was even more pronounced when combined, with a noticeable impact already apparent at 4 hours (Figure 4A-B). The combination also exhibited a decreasing trend in total PLCγ2 levels at 24 and 48 hours, consistent with the RNA-Seq data (Figure 3C). Furthermore, it significantly inhibited PLCγ2-Y1217 phosphorylation more effectively than zanubrutinib alone, particularly at 48 hours (Figure 4A-B). Moreover, p65 protein levels were downregulated considerably only by the combination treatment at 24 hours, whereas none of the single agents had this effect. The phosphorylation of p65 on serine 536 was reduced with zanubrutinib alone and further suppressed by the combination with rigosertib (Figure 4A-B).

**Figure 4.**
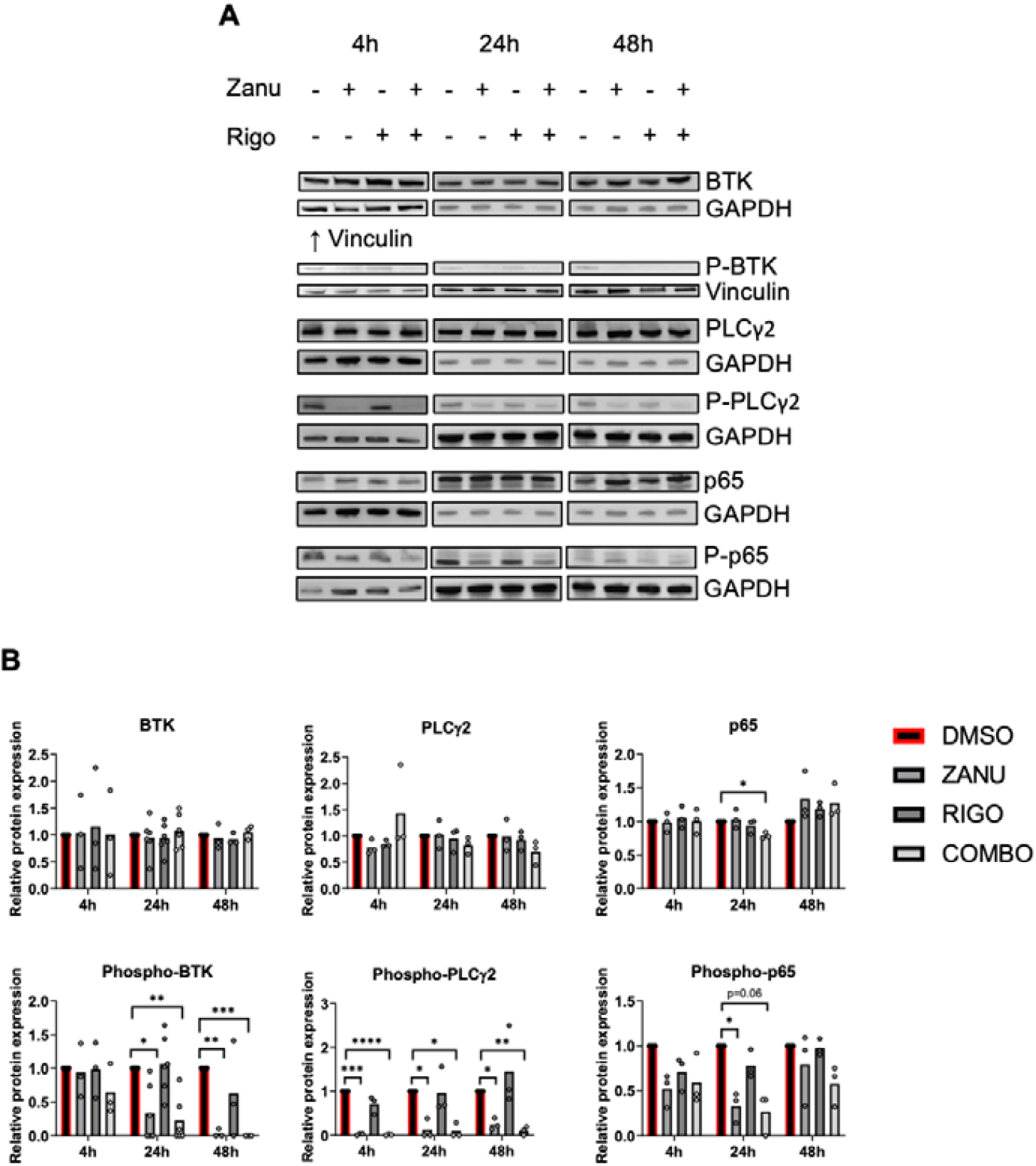
Rigosertib synergizes with zanubrutinib in inhibiting the phosphorylation of BTK, PLCγ2, and p65 on activation sites. A) BTK, phospho-BTK (Y223), PLCγ2, phospho-PLCγ2 (1217), p65 and phospho-p65 (S536) protein levels in Karpas1718 parental cells upon zanubrutinib (5 nM), rigosertib (10 nM) or zanubrutinib/rigosertib combination. GAPDH or vinculin was used as a loading control. At least three biological replicates are shown, with one representative replicate illustrated. B) BTK, phospho-BTK (Y223), PLCγ2, phospho-PLCγ2 (1217), p65, and phospho-p65 (S536) protein levels in Karpas1718 parental cells upon zanubrutinib (5 nM), rigosertib (10 nM), or zanubrutinib/rigosertib combination, assessed by densitometric quantification. Each dot represents a biological replicate normalized to the respective loading control. Results coming from at least three biological replicates. Statistical significance tested with Two-way ANOVA + Dunnett’s multiple comparisons test (* = p < 0.05, ** = p < 0.01, *** = p < 0.001). (ZANU=zanubrutinb; RIGO=rigosertib; COMBO=combination).

In the canonical NF-κB signaling pathway, the transcription factor p65 is downstream of BCR signaling activation, BTK, and PLCγ2 phosphorylation. Upon phosphorylation, p65 accumulates in the nucleus and activates the NF-κB transcriptional program ^9^. To better understand the role of rigosertib in BCR signaling and NF-κB activation, we investigated p65 localization by immunofluorescence upon treatment with zanubrutinb (5 nM), rigosertib (10 nM), and the combination at 4, 6, and 8 hours. The combination of zanubrutinb and rigosertib significantly reduced p65 nuclear translocation at 8 hours (Figure 5A-B) without affecting total p65 protein levels at the analyzed time points (Figure 5C), in accordance with the immunoblot results (Figure 4B).

**Figure 5.**
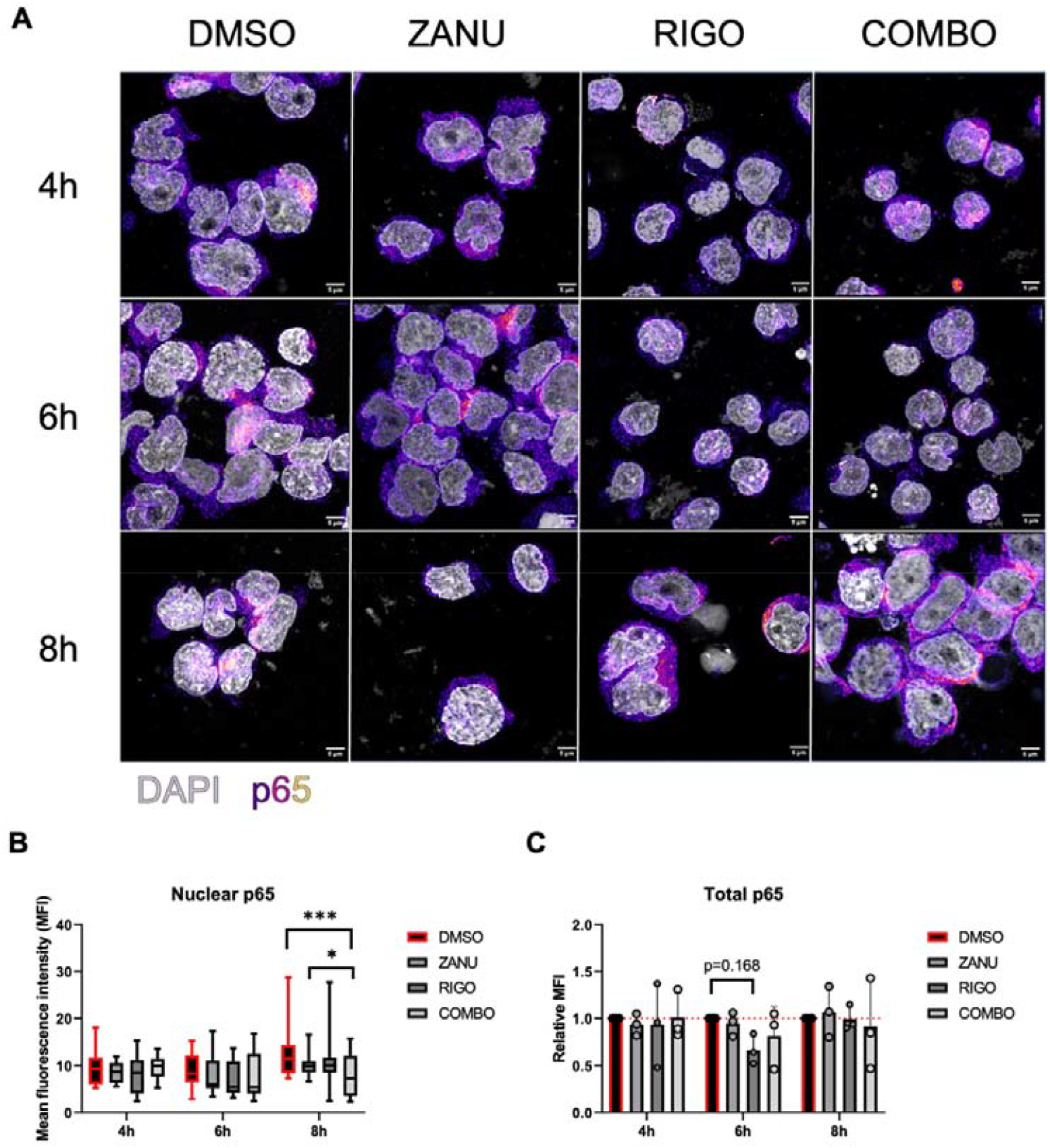
p65 nuclear protein levels are reduced upon zanubrutinib and rigosertib combination. A) Representative immunofluorescence maximum projection images of Karpas1718 parental cells upon zanubrutinib (5 nM), rigosertib (10 nM) or zanubrutinib/rigosertib combination. Nuclei in gray, p65 signal, Fire lookup table colored (Fiji). 3X zoom of images taken with Leica TCS SP5 confocal_microscope with HCX PL APO lambda blue 63.0 × 1.40 OIL objective. Scale bar = 5 μm. B) Quantification of nuclear p65 on maximum projection images of Karpas1718 parental cells upon zanubrutinib (5 nM), rigosertib (10 nM), or zanubrutinib/rigosertib combination. Results coming from three biological replicates. Statistical significance tested with Two-way ANOVA + Dunnett’s multiple comparisons test (* = p < 0.05, ** = p < 0.01, *** = p < 0.001). C) Quantification of total p65 on maximum projection images of Karpas1718 parental cells upon zanubrutinib (5 nM), rigosertib (10 nM), or zanubrutinib/rigosertib combination. Each dot represents a biological replicate normalized to the DMSO control. Results coming from three biological replicates. Statistical significance tested with Two-way ANOVA + Dunnett’s multiple comparisons test (* = p < 0.05, ** = p < 0.01, *** = p < 0.001).

We evaluated whether rigosertib had any effect on PI3K/AKT/mTOR signaling in our model to rule out the possibility that the reduction in BCR signaling is related to rigosertib off-target effects. At the transcript level, 10 nM rigosertib had no significant effects on the PI3K/AKT/mTOR signaling pathway in Karpas1718 parental cells (Supplementary Figure 4A). Consistently, 10nM rigosertib had no effects on total AKT protein levels and phosphorylation of AKT on serine 473 at 24 hours in Karpas1718 parental cells, as shown by western blot (Supplementary Figure 4B). In addition, to assess whether rigosertib might influence the kinetics of AKT activation, Karpas1718 parental cells were transfected with the FRET-based AKT activity reporter Lyn-AktAR2-EV or a control “dead” unresponsive version of the reporter (Lyn-AktAR2-EV-D) as previously described ^37,38^. Active and control reporter transfected K1718 cells were incubated for 16 hours with either 10 nM rigosertib or DMSO as a control. Afterwards, anti-IgM or anti-IgG stimulation was used to trigger AKT activation, and AKT activity over time was measured by flow cytometry in a kinetics setting. As expected, anti-IgM but not anti-IgG stimulation led to an extensive activation of AKT in K1718 parental cells transfected with the AKT FRET-reporter but not in the dead reporter-transfected cells. Pre-treatment with rigosertib did not affect AKT activation kinetics in Karpas1718 (Supplementary Figure 4C). Moreover, we did not observe significant effects on AKT activity, as measured by flow cytometry, upon 4-hour exposure to increasing rigosertib concentrations in Karpas1718-EV (Supplementary Figure 4D). Only when rigosertib was combined with zanubrutinib did we observe downregulation at the transcript level of the PI3K/AKT/mTOR signaling pathway (Supplementary Figure 4A) and, consistently, decreased AKT activity at several concentrations (Supplementary Figure 4D). This suggests that the combination may have a synergistic effect on inhibiting PI3K/AKT/mTOR signaling.

In summary, rigosertib synergizes with zanubrutinib by enhancing the inhibition of the BCR signaling through the canonical NF-κB pathway.

## Discussion

In this study, a screen of 1,695 compounds in the bona fide MZL model Karpas1718 and its derivative with secondary resistance to BTK and PI3K inhibitors identified 33 small molecules with high single-agent activity (IC50 < 100 nM) in both cell types. From these, we selected candidates based on their clinical potential, validating molecules targeting PLK1 (rigosertib), AURKA (alisertib), HSP90 (ganetespib), HDAC (fimepinostat, panobinostat), and metabolic processes (elesclomol, disulfiram) for their ability to enhance the anti-tumor activity of BTK and PI3K inhibitors in multiple MZL cellular models. PLK1 is a key cell cycle regulator, involved in various stages of mitosis, particularly at the mitotic entry stage, and is frequently overexpressed in cancer ^42,44-46^. PLK1 inhibitors have demonstrated preclinical anti-tumor activity in several hematological and solid tumors, although regulatory agencies have yet to approve any compound ^47-52^. We focused on rigosertib ^47^, a representative of the PLK1 inhibitors, due to its potent activity and ability to overcome resistance to BTK inhibitors in our models. Moreover, it has demonstrated activity in additional preclinical models, also derived from chronic lymphocytic leukemia and diffuse large B-cell lymphoma, and has an acceptable safety profile, although with limited activity as a single agent in the clinical setting ^51-55^. We found that the combination of rigosertib with the second-generation BTK inhibitor zanubrutinib in Karpas 1718 parental cells led to the downregulation of key biological processes, including nuclear transport, RNA processing, and immune-related pathways. Notably, among the immune-related pathways, those involved in BCR signaling and B-cell activation and proliferation were significantly suppressed by the combination treatment. To validate these findings, we examined total and phosphorylated levels of three key proteins of the BCR signaling axis: BTK, PLCγ2, and p65. Rigosertib, combined with zanubrutinib, further reduced phosphorylation at their activating sites: BTK-Y223, PLCγ2-Y1217, and p65-S536 ^8,56^.

While the combination treatment did not have a significant effect on total protein levels, it reduced the phosphorylation and nuclear accumulation of p65, thereby inhibiting NF-κB activation. This supports the additive inhibitory effect of zanubrutinib and rigosertib on BCR signaling in MZL.

Our study presents the first evidence demonstrating that rigosertib synergizes with BTK inhibitors in B-cell lymphoma by enhancing BCR inhibition, primarily due to reduced activation of the canonical NF-κB pathway. Rigosertib has been described as inhibiting the PI3K/AKT/mTOR pathway in head and neck squamous cell carcinoma ^57^ and myelodysplastic syndrome^58^ models. We demonstrated that rigosertib alone did not affect PI3K/AKT/mTOR signaling at the transcript and protein levels, nor did it alter AKT activity in our model. However, we observed that the combination of rigosertib and zanubrutinib may affect PI3K/AKT/mTOR signaling, though this observation will require further investigation.

We did not directly investigate the activity of zanubrutinib and rigosertib on the cell cycle. However, we showed that rigosertib induced the upregulation of essential genes involved in microtubules and mitotic spindle assembly, confirming PLK1 targeting in our model. PLK1 inhibitors have already been reported to arrest the cell cycle in other B-cell lymphoma models, including MZL ^50^ and mantle cell lymphoma (MCL) ^59^. Jiang et al. showed that combining acalabrutinib, a second-generation BTKi, with volasertib, a second-generation PLK1 inhibitor, causes cell cycle arrest and exhibits synergistic effects in MCL models. However, the combination did not enhance cell cycle arrest beyond that of volasertib alone, suggesting that other mechanisms, such as BCR signaling inhibition, may underlie the observed synergy ^59^.

PLK1 has emerged as a promising therapeutic target due to its role in mitosis and its frequent overexpression in various types of cancer ^60^. Moreover, PLK1 inhibition can overcome resistance to the BCL2 inhibitor venetoclax in preclinical models of T-cell acute lymphoblastic leukemia ^61^.

Consistent with these findings, our results suggest that PLK1 inhibition holds strong clinical potential for treating MZL, with potential to be extended in other hematological malignancies. Combination therapies provide an effective strategy to overcome therapeutic resistance and minimize residual disease. In conclusion, our study identified several compounds with potent anti-tumor activity in MZL models, which was maintained in a model of secondary resistance to BTK inhibition. This provided the first evidence that rigosertib can overcome resistance to BTK inhibitors in MZL.

## Acknowledgments

The authors used OpenAI’s ChatGPT (GPT-5; OpenAI, San Francisco, CA, USA) to assist in text preparation and drafting. All content was critically reviewed, revised, and verified by the authors, who take full responsibility for the final manuscript.

## Author contribution

AZ: performed experiments, analyzed and interpreted data, performed data mining, prepared the figures, and co-wrote the original draft.

AJA: performed experiments, analyzed and interpreted data, performed data mining, prepared the figures, and co-wrote the original draft.

MR: performed experiments, analyzed, and interpreted data. LC: performed data mining.

SV, AA, GV, KK, EC, FS, GS, CT, SN performed experiments. MM, OH: provided advice and resources.

EZ, DR, AS: provided advice.

AR: performed genomics experiments.

MV, CM: co-designed research and interpreted data.

FB: co-designed research, interpreted data, and co-wrote the original draft.

All authors reviewed and accepted the final version of the manuscript.

