## Supplementary tables and figures for "High-Throughput Screening Identifies PLK1 Inhibition as a Strategy to Potentiate BTK Blockade in Marginal Zone Lymphoma"

*[1] Institute of Oncology Research, Faculty of Biomedical Sciences, USI, Bellinzona, Switzerland; [2] Regenerative Medicine Division, Institute for Translational Research, Università della Svizzera Italiana (USI) - Ente Ospedaliero Cantonale (EOC), Bellinzona, Switzerland; [3] Swiss Institute of Bioinformatics, Lausanne, Switzerland; [4] Experimental Therapeutics Program, IFOM ETS - The AIRC Institute of Molecular Oncology Milan, Italy; [5] Biocev, First Faculty of Medicine, Charles University, Prague, Czech Republic; [6] Oncology Institute of Southern Switzerland, Ente Ospedaliero Cantonale (EOC), Bellinzona, Switzerland; [7] Department of Hematology, First Faculty of Medicine, Charles University and General University Hospital, Prague, Czech Republic.*

*Equally contributed

**Corresponding author**: Prof. Francesco Bertoni, Institute of Oncology Research, Faculty of Biomedical Sciences, USI, Bellinzona, via Francesco Chiesa 5, 6500 Bellinzona, Switzerland. Phone: +41 58 666 7206;

### Supplementary Materials and Methods

#### Supplementary Tables

**Supplementary Table 1.** Antibodies/molecules used for immunoblotting (IB) or immunofluorescence (IF) as indicated in the Materials and Methods section.

| Antibody | Catalog Number | Company | Type | Host | Dilution Used |
|---|---|---|---|---|---|
| BTK | 8547S | Cell Signaling | Primary | Rabbit | 1:1000 (IB) |
| Phospho-BTK (Y223) (D9T6H) | 87141S | Cell Signaling | Primary | Rabbit | 1:1000 (IB) |
| BCL-2 (C-2) | sc-7382 | Santa Cruz Biotechnology | Primary | Mouse | 1:1000 (IB) |
| PLCγ2 (E5UKT) | 55512s | Cell Signaling | Primary | Rabbit | 1:1000 (IB) |
| Phospho-PLCγ2 (Tyr1217) (E2U1K) | 54442s | Cell Signaling | Primary | Rabbit | 1:1000 (IB) |
| NF-kappaB p65 (C22B4) | 4764S | Cell Signaling | Primary | Rabbit | 1:1000 (IB); 1:200 (IF) |
| P-NF-kappaB p65 (S536) (93H1) | 3033S | Cell Signaling | Primary | Rabbit | 1:1000 (IB) |
| AKT | 9272S | Cell Signaling | Primary | Rabbit | 1:1000 (IB) |
| P-AKT (S473) (D9E) XP(R) | 4060L | Cell Signaling | Primary | Rabbit | 1:1000 (IB) |
| Vinculin | V9131 | Merck | Primary | Mouse | 1:1000 (IB) |
| GAPDH | FF26A | Invitrogen | Primary | Mouse | 1:1000 (IB) |
| Phalloidin iFluor 633 | ab176758 | Abcam | | | 1:1000 (IF) |
| Anti-Rabbit Alexa Fluor 488 | A-11008 | | Secondary | Goat | 1:1000 (IF) |

**Supplementary Table 2.** Compounds screened in the library and their effect on Karpas1718 parental and resistant cells.

#### Supplementary Figures

**Supplementary Figure 1.** A) Dose-response curves in parental (PAR) and resistant (IDE) Karpas1718 and VL51 cells treated with increasing idelalisib concentration and fixed disulfiram concentration (0.25µM). Graphs generated with R software. B) Dose-response curves in parental (PAR) and resistant (IDE/IBR) Karpas1718 and VL51 cells treated with increasing ibrutinib concentration and fixed disulfiram concentration (0.25µM). Graphs generated with R software.

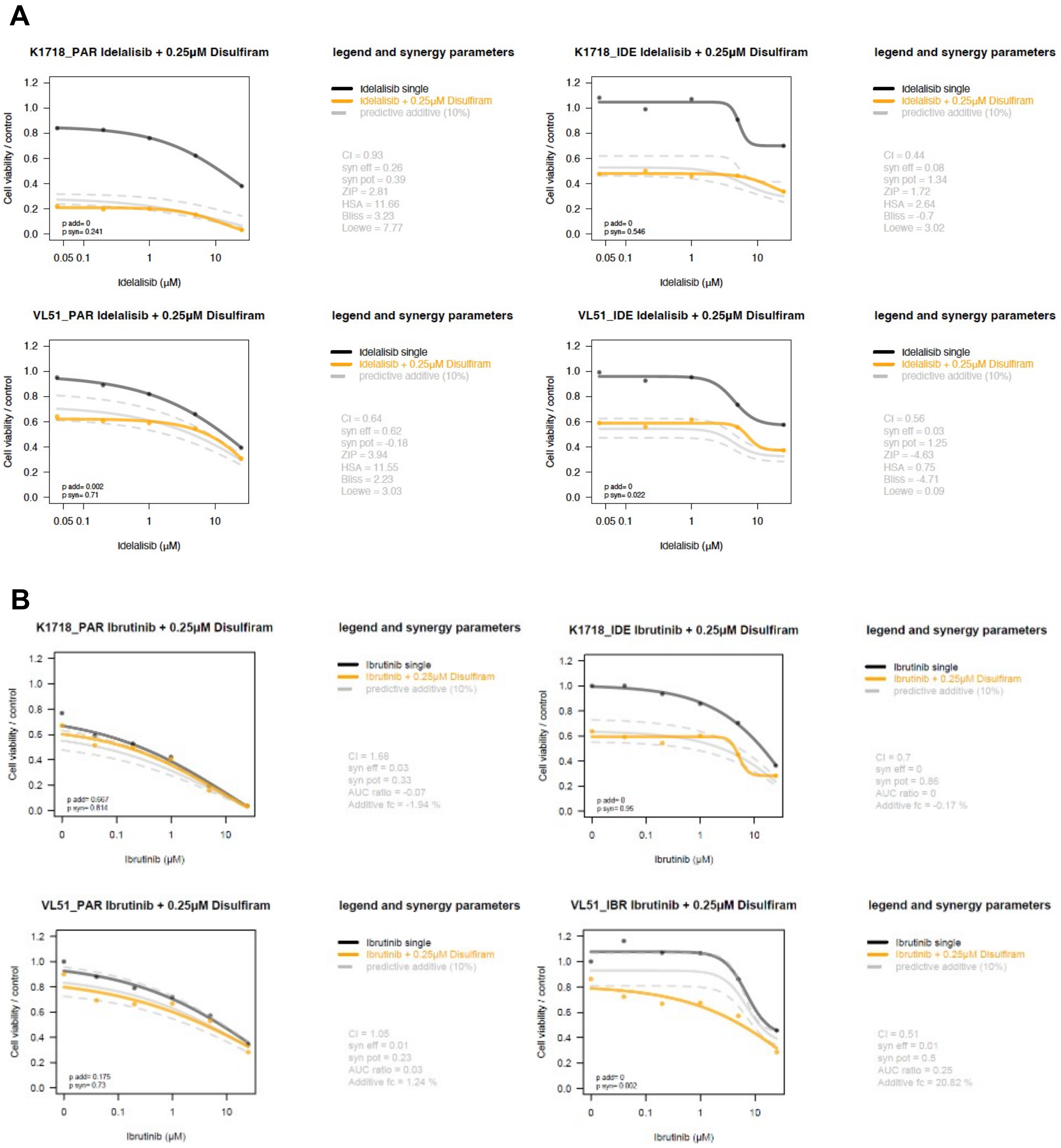

**Supplementary Figure 2.** A) Enrichment plots of the indicated pathway upon zanubrutinib (5nM), rigosertib (10nM), or zanubrutinib/rigosertib combination in Karpas1718 parental cells. Graphs generated with R software. B) Ridge plots on top 30 GOCC pathways upon zanubrutinib (5nM), rigosertib (10nM), or zanubrutinib/rigosertib combination in Karpas1718 parental cells. Graphs generated with R software. C) Enrichment plots of the indicated pathway upon zanubrutinib (5nM), rigosertib (10nM), or zanubrutinib/rigosertib combination in Karpas1718 parental cells. Heatmap showing the regulation of the core enrichment genes upon the indicated treatment conditions. Graph generated with R software. Statistical significance tested with moderated t-test (*=p<0.05; **=p<0.01; ***=p<0.01). (ZANU=zanubrutinb; RIGO=rigosertib; COMBO=combination)

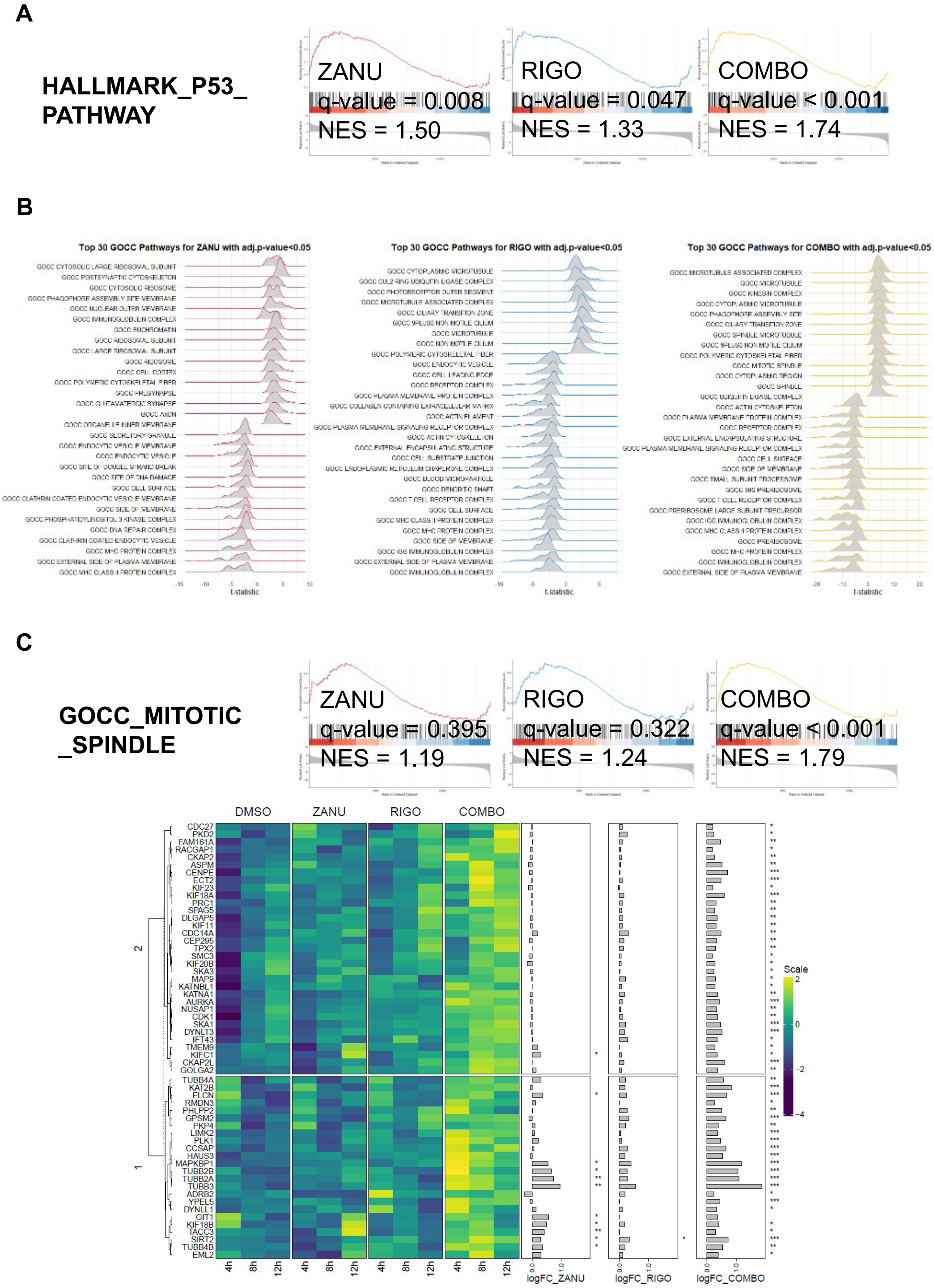

**Supplementary Figure 3.** A) Enrichment plots of the indicated pathway upon zanubrutinib (5nM), rigosertib (10nM), or zanubrutinib/rigosertib combination in Karpas1718 parental cells. Graphs generated with R software. B) Heatmap showing the regulation of the core enrichment genes upon the indicated treatment conditions. Graph generated with R software. Statistical significance tested with moderated t-test (*=p<0.05; **=p<0.01; ***=p<0.01). C) BCL-2 protein levels in Karpas1718 parental cells upon zanubrutinib (5nM), rigosertib (10nM), or zanubrutinib/rigosertib combination. GAPDH or vinculin were used as loading controls. At least three biological replicates are shown, with one representative replicate illustrated. D) BCL-2 protein levels in Karpas1718 parental cells upon zanubrutinib (5nM), rigosertib (10nM), or zanubrutinib/rigosertib combination assessed by densitometric quantification. Each dot represents a biological replicate normalized to the respective loading control. Results coming from at least three biological replicates. Statistical significance was assessed using a two-way ANOVA followed by Dunnett's multiple comparisons test.

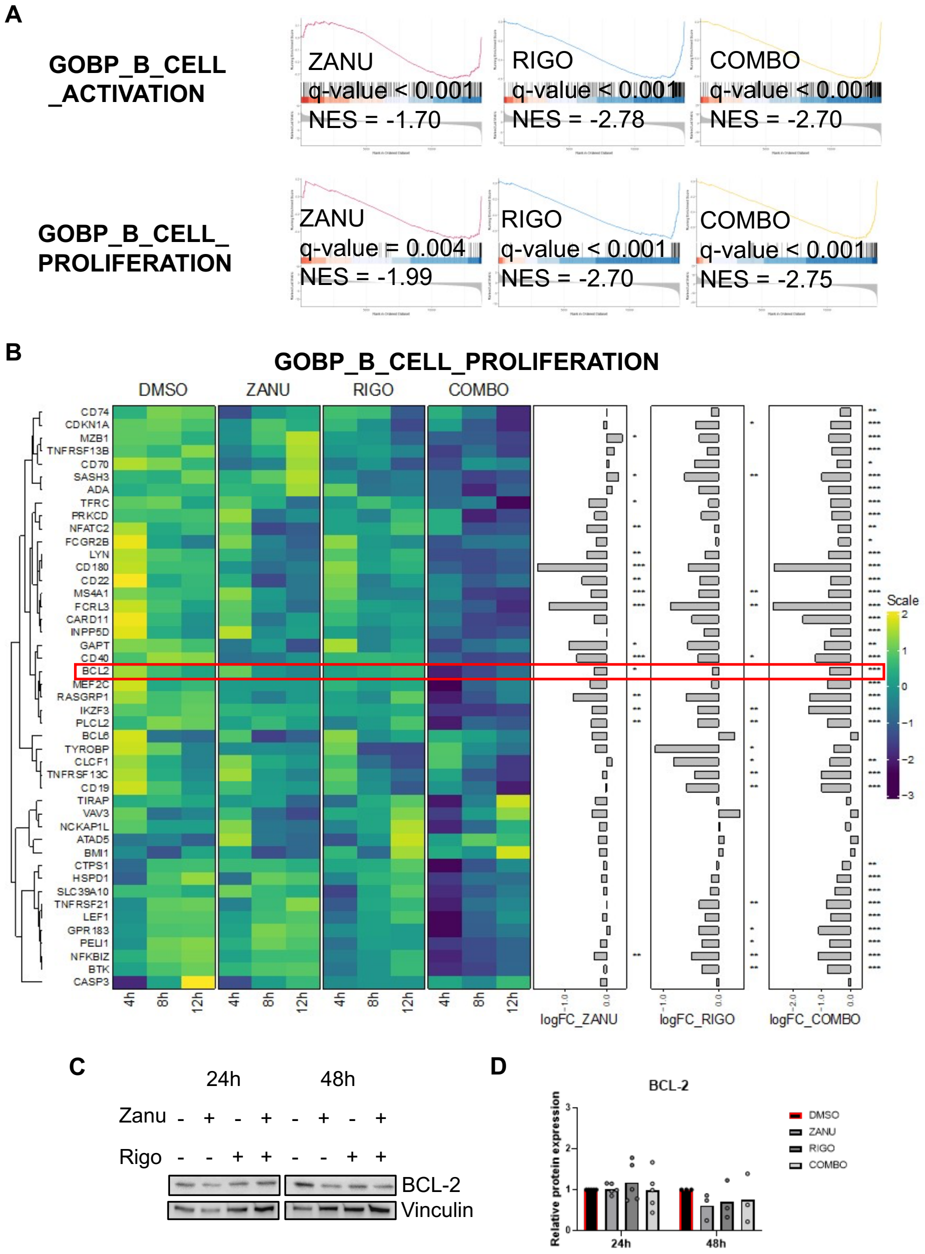

**Supplementary Figure 4.** A) Enrichment plots of the indicated pathway upon zanubrutinib (5nM), rigosertib (10nM), or zanubrutinib/rigosertib combination in Karpas1718 parental cells. Graphs generated with R software. B) AKT and phospho-AKT (S473) protein levels in Karpas1718 parental cells upon zanubrutinib (5nM), rigosertib (10nM), or zanubrutinib/rigosertib combination. Vinculin was used as a loading control. One biological replicate. C) Kinetics of AKT activity upon rigosertib (10nM) or DMSO after BCR stimulation with anti-IgG or anti-IgM (5 µg/mL) stimulation measured by FRET. Values shown are normalized to the baseline FRET signal of Karpas1718 before antigenic stimulation. D) AKT activity upon increasing concentration of zanubrutinib, rigosertib, or zanubrutinib/rigosertib combination measured by FRET. The values shown were first normalized to the baseline FRET signal of Karpas1718 transfected with a dead, unresponsive reporter to account for nonspecific changes in FRET, and then normalized to the baseline FRET without any inhibitor.

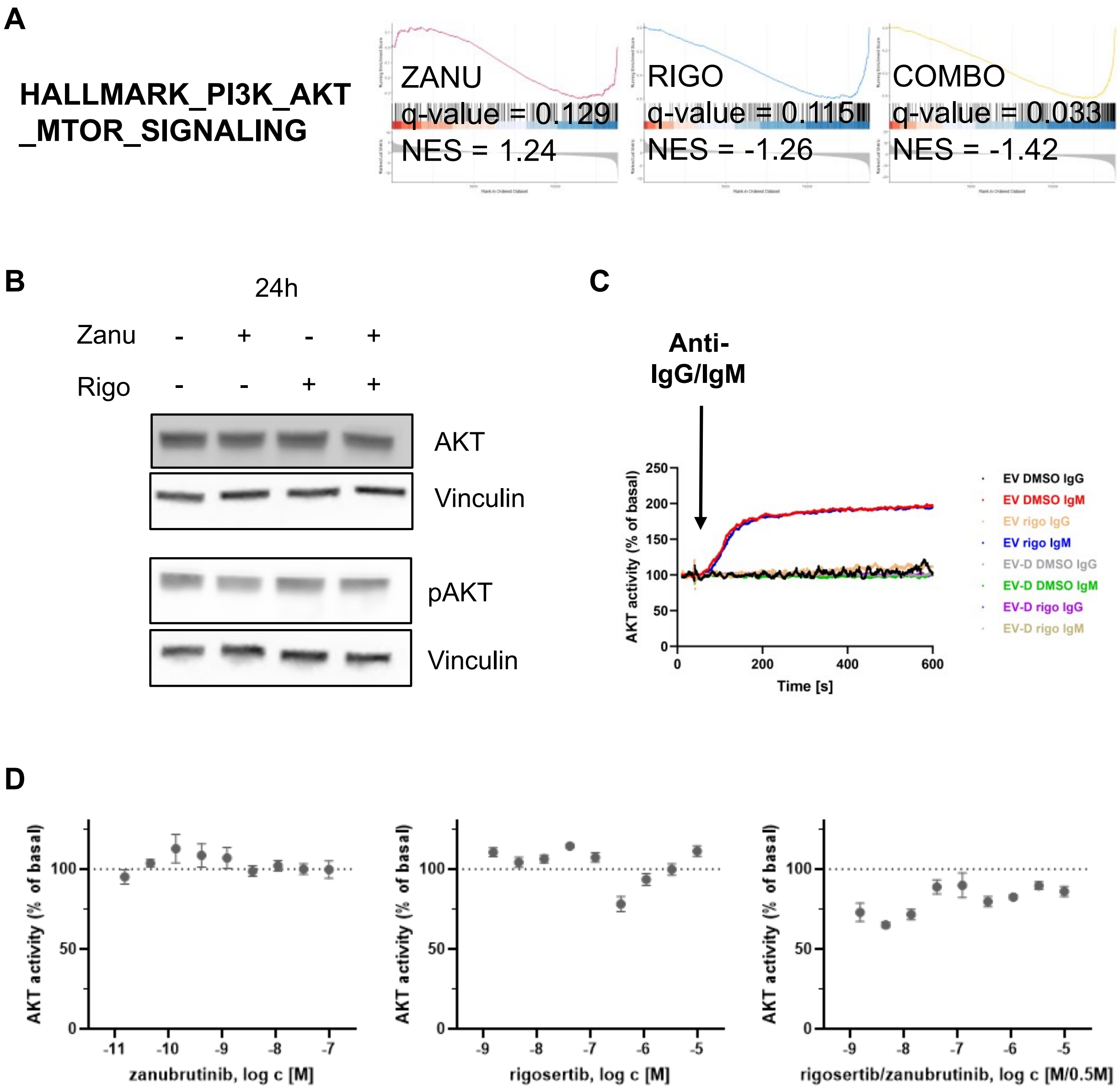
